# Explainable machine learning for the identification of proteome states via the data processing kitchen sink

**DOI:** 10.1101/2023.08.30.555506

**Authors:** Aaron M. Scott, Erik Hartman, Johan Malmström, Lars Malmström

## Abstract

The application of machine learning algorithms to facilitate the understanding of changes in proteome states has emerged as a promising methodology in proteomics research. Unfortunately, these methods can prove difficult to interpret, as it may not be immediately obvious how models reach their predictions. We present the data processing kitchen sink (DPKS) which provides reproducible access to classic statistical methods and advanced explainable machine learning algorithms to build highly accurate and fully interpretable predictive models. In DPKS, explainable machine learning methods are used to calculate the importance of each protein towards the prediction of a model for a particular proteome state. The calculated importance of each protein can enable the identification of proteins that drive phenotypic change in a data-driven manner while classic techniques rely on arbitrary cutoffs that may exclude important features from consideration. DPKS is a free and open source Python package available at https://github.com/InfectionMedicineProteomics/DPKS.

## 1 Introduction

Developments of proteomic data acquisition technologies have resulted in a significant increase in throughput, facilitating the expansion of proteomics experiments from 10s to 1000s of samples. The detailed proteome maps produced require multiple downstream tools to get to interpretable biological results from the raw data as there remains a lack of coherent and standardized pipelines to conduct the analysis. Robust tools to apply normalization [1], protein quantification [2–5], differential expression [6–10], and machine learning algorithms to the results are not all readily available to those without advanced programming expertise. State-of-the-art tools must be selected and placed logically together in a pipeline in order to glean biological insight from raw proteome maps, which is not a trivial endeavor. Although there are some tools that expose portions of the pipeline and attempt to present the complex algorithms using a graphical user interface to non-programmers, they remain closed-source and restrictive in their usage [11]. The open-source alternatives are typically implemented in the R programming language and require different file formats as input, requiring programming to sequentially apply different tools, making analytical experimentation and evaluation more difficult. Additionally, many of these tools do not follow standard software design principles, making the code difficult to understand and tedious to extend, even if it is open-source. The variety of analytical options and lack of standardization hinders the reproducibility of analysis and could benefit from consistently available methods that adhere to the same standard.

Recently, the application of machine learning to proteomics experiments has been shown to improve the identification and selection of prognostic protein biomarkers in disease [12, 13]. However, the implementation of machine learning algorithms in clinical protein biomarker detection experiments remains non-trivial, and expert knowledge must be used to select the correct hyperparameters, deal with class imbalances, and correctly deal with low sample numbers to ensure the models are trained properly and provide results that can generalize to new data. Additionally, many complex models are difficult to interpret, as it may not be immediately obvious how models reach their predictions. To fully understand the behavior of a model, explainable machine learning and feature attribution methods, such as SHAP [14], are used to determine the importance of each input protein to the predictions of a model. These feature attribution methods result in fully explainable machine learning models, where the importance of each input is explicitly calculated. The calculated importance for each protein can facilitate the selection of potential prognostic protein biomarkers in a fully data-driven approach, while classic techniques rely on arbitrary cutoffs that may exclude important features from further analysis. Currently, there is no tool available to combine all of these steps to produce robust and explainable machine learning models for proteomics data, so expert knowledge of programming is required to implement them.

We present the data processing kitchen sink (DPKS) that combines multiple analytical steps into a singular Python package, and exposes them via an expressive and user-friendly application programming interface (API) for the in-depth analysis of proteomic data. DPKS provides easy and efficient access to state-of-the-art statistical analysis algorithms for normalization, protein quantification, and differential expression and provides the ability to train and interpret complex machine learning models with a single line of code. Using the explainable machine learning methods available via DPKS, it is possible to select highly accurate panels of proteins for classifying disease that could be missed using classic statistical analysis. We verify and display the capabilities of DPKS using a spike-in 2-species proteome mixture experiment, and then apply the DPKS package to find a panel of highly predictive protein biomarkers to identify severe COVID-19 [15] in minutes on a standard laptop with no specialized hardware. Additionally, we highlight how these proteins may be missed for selection during classic statistical analysis, and how a combination of reproducible statistical methods in combination with explainable machine learning can guide the investigation of biological systems and expedite the identification of potential prognostic protein panels in an automated and data-driven manner. DPKS is open-source and available for free at https://github.com/InfectionMedicineProteomics/DPKS.

## 2 Results

DPKS comprises a suite of statistical processing algorithms for the in depth analysis of proteomics data. Encompassing every step of the downstream analysis pipeline, DPKS provides access via our user-friendly API to 5 main analytical steps (**Figure 1**). First, we provide parsers for the direct input from a number of common mass spectrometry signal processing tools, as well as a simple tabular format that is easy to conform to. Second, we provide access to several sample normalization methods, such as total ion chromatogram, mean, and median sample normalization, and the option to perform retention time window normalization [1]. Third, we provide multiple quantification algorithms, such as top-N for absolute quantification, and the iq method for relative quantification [3] to calculate protein abundances from precursors. These quantification algorithms can be used at multiple levels to directly quantify precursors, peptides, or proteins. Forth, we provide multiple statistical tests for experiment-wide differential abundance experiments using t-tests, linear regression, or ANOVA, and a variety of multiple testing correction algorithms to control the false discovery rate. Fifth, we provide easy access to advanced machine learning algorithms for model selection and hyper-parameter optimization, model training via cross-validation, feature attribution using SHAP [14], as well as complete feature ranking and biomarker panel selection using recursive feature elimination. Additionally, we expose important methods for missing value imputation as well as plotting functions to easily visualize data.

**Figure 1:**
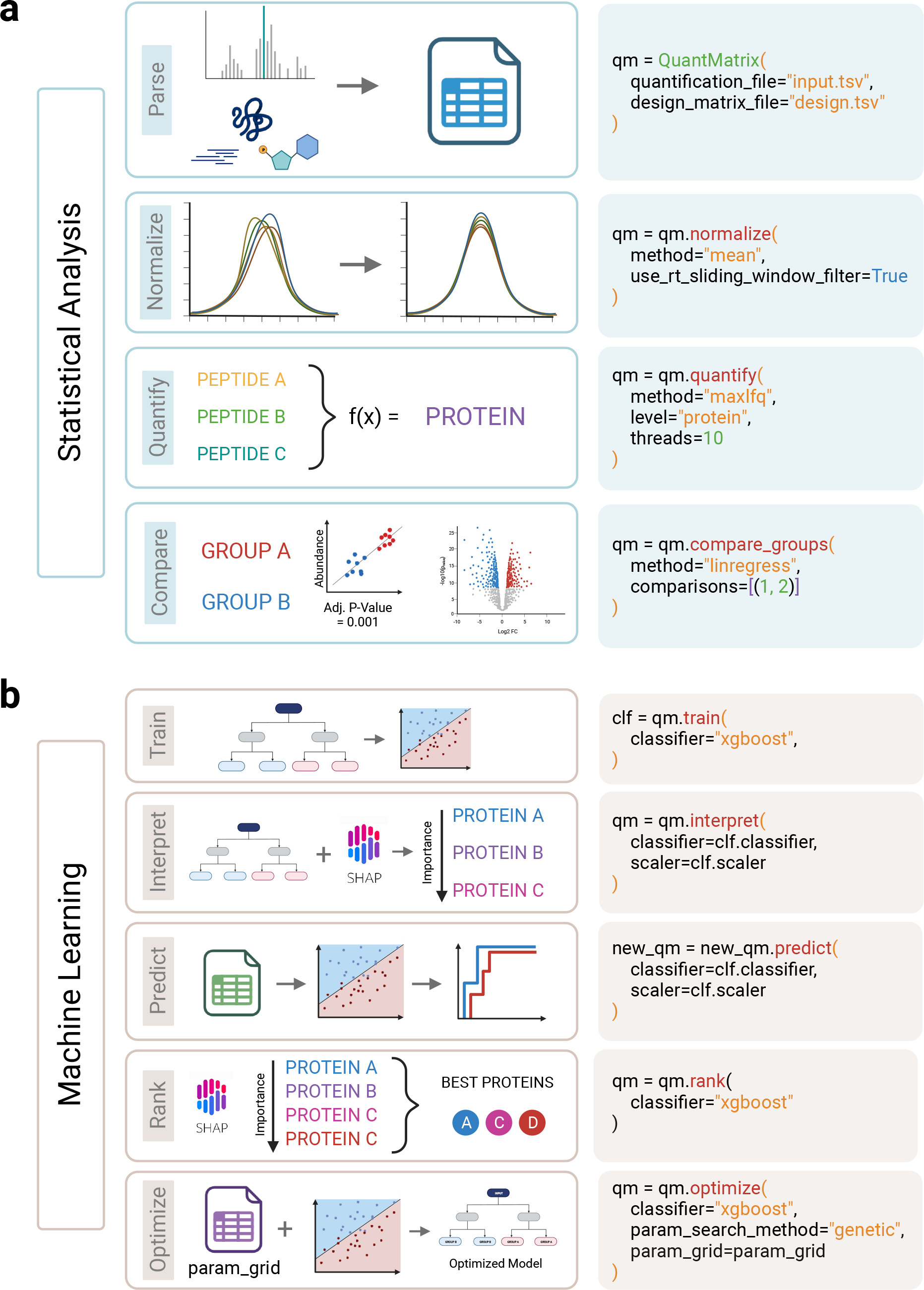
Overview of the DPKS pipeline and software package. **a** Shows the functionality in DPKS related to statistical analysis, specifically to perform differential abdunance analysis in a reproducible manner. The functions, and how you call them using the Python package are shown step-by-step. **b** Shows the functionality in DPKS related to machine learning and how to rank and identify the most predictive proteins for a comparison. (Created with BioRender.com)

Each one of the above functions can be called using one line of code, and data frames can be exported whenever during the analytical process. By utilizing method-chaining in the main data structure of DPKS we can build modular and complex workflows for customized analysis in a simple manner. To validate that each step of the workflow is performing correctly, we first analyze the normalization, quantification, and differential expression functionality of DPKS using samples containing tryptic yeast peptides spiked in to a constant mouse-kidney proteome tryptic digest in a dilution series resulting in 6 distinct groups with 10 samples per group at known concentrations of yeast peptides [12]. Since we know the expected ratios of yeast and mouse proteins between sample groups, we can use this data to verify that the algorithms implemented in DPKS are working correctly to quantify proteins. Previously, we have used DPKS to select a panel of protein biomarkers for septic AKI [12] and applied similar feature attribution methods to provide high accuracy classifiers and intelligent pathway analysis using biologically informed neural networks (BINN) [13] to show how protein biomarker panels can be identified using explainable machine learning that outperform proteins selected via classic statistical analysis. Here, we refine and present our previous methods in a coherent package to reanalyze a dataset consisting of patients with COVID-19 at varying levels of severity (1-7 WHO-grade)[15] using the full suite of DPKS methods. As we have previously reanalyzed this data to demonstrate the application of BINNs [13], here, we additionally show that explainable machine learning with recursive feature elimination can be used to identify a panel of 10 highly accurate predictive protein biomarkers in minutes that may be overlooked using classic statistical analysis.

### 2.1 DPKS provides easy access to common statistical methods

Normalization between samples serves to minimize technical variation, correct batch effects, and ensure data from different samples are placed on the same scale so that it is comparable. We show the effects of normalization on a serial dilution (spike-in) dataset, where increasing amounts (1X, 2X, 4X, 8X, 16X, 32X) of yeast peptides were spiked-in to a constant background mouse proteome [12]. This will allow for the direct evaluation of the methods in DPKS based on the measurements of known yeast ratios between steps in the dilution series. The unnormalized intensity distributions for each sample show marked variability between groups (**Figure 2a**), while normalization transforms the abundance distributions so that all are on the same scale and can be compared to one another (**Figure 2b**). Additionally, density plots for the abundance profile of each unnormalized sample group distinctly in batches based on their dilution series factor (**Figure 2c**) making comparisons between groups difficult, as technical variation between groups may be misinterpreted as biological signal. Normalization adjusts the means and overall distributions of the abundance profiles so they are overlapping, there by removing the distinct grouping for each sample (**Figure 2d**). We further analyze the effect of normalization using principal component analysis (PCA) (**Figure 2c-d**). The silhouette score, which measures how similar an instance is to their assigned cluster compared to other clusters, for unnormalized data is 0.61 (**Figure 2e**) compared to 0.86 for normalized data (**Figure 2f**). This 40.98% increase shows that the individual groups cluster more accurately together after the technical variation in an experiment is removed using normalization. DPKS provides a fast and easy to use interface to access normalization methods with just one function call (**Figure 1a**).

**Figure 2:**
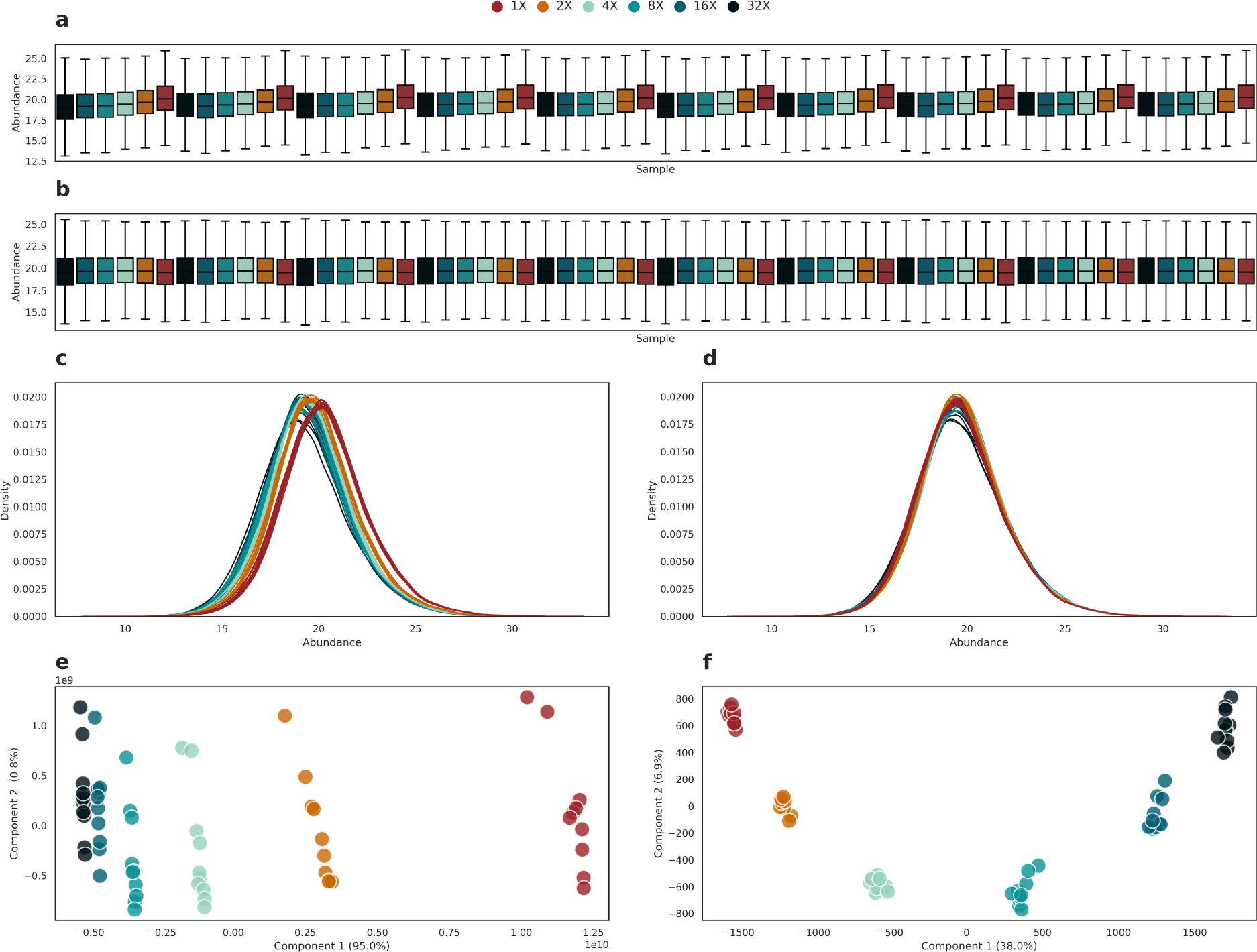
The effects of between-sample normalization. Normalization reduced technical variability between samples and reduces batch effects between sample groups to allow for accurate comparisons. **a** shows unnormalized boxplots for each sample the in spike-in data colored by dilution factor. **b** shows the same samples after normalization. Here, the variability between sample groups is minimized and the means between all samples are on the same level. **c** shows density plots of the abundance for each sample colored by dilution factor. **d** shows the sample samples after normalization. This shows a similar effect to **b** as the batches between abundance profiles are removed after normalization. **e** shows the PCA of the spike-in samples colored by dilution. **f** shows the sample samples after normalization and a marked increase in the tightness of the displayed clusters.

To measure the protein content of a sample using mass spectrometry proteomics, proteins are typically digested using trypsin into peptides for downstream analysis. Protein quantities are then inferred from peptide quantities, which is a necessary and non-trivial step in the analysis of proteomics data. Numerous methods have been developed to quantify proteins, and DPKS currently provides access to both absolute quantification (topN), and relative quantification (MaxLFQ [4], iq [3]) methods. It should be clarified that methods such as iq [3] and MaxLFQ [4] are relative quantification methods, meaning that they provide optimal protein abundances for comparing between groups, while absolute quantification methods (topN) provide a means to measure the absolute abundance of proteins in a proteome. DPKS supports the quantification of features at the precursor and peptide level using these methods, although here we focus on demonstrating the effects of relative quantification via the DPKS implementation of the iq algorithm [3] (**Figure 3a-b**). Intensities between 2 groups of the spike-in data, 4X vs 16X, were compared at the precursor and protein level (**Figure 3a-b**). Here, we expect a 4-fold change (2 on a log2 scale) of yeast identifications and a constant level of mouse identifications (0 on a log2 scale) between measured groups. The noise in the data between groups (**Figure 3a**) is significantly decreased after relative protein quantification while maintaining fold changes around the expected ratios (**Figure 3b**), demonstrating how biological signal can be preserved while providing more meaningful annotations to the data (i.e. proteins instead of peptides or precursors). Dotted lines indicate the expected ratios and solid lines around the expected ratios visualize ± 0.2 windows that are used to define the ratio validated identification rates [16]. Within the ratio validated regions, the combined false discovery rate, defined as mouse identifications within the 4-fold change region or yeast identifications within the 0-fold change region, is only 3.3%. The total recall of differentially abundant proteins, defined as mouse proteins within the 0-fold change region combined with yeast proteins within the 4-fold change region is 65.1%. The lower recall of total proteins may be attributed here to missing values in the 16X dilution samples due to lower abundance. Via the expressive API, DPKS supports easy access to different methods of quantification (**Figure 1a**) allowing for the optimal method to be evaluated for a given dataset.

**Figure 3:**
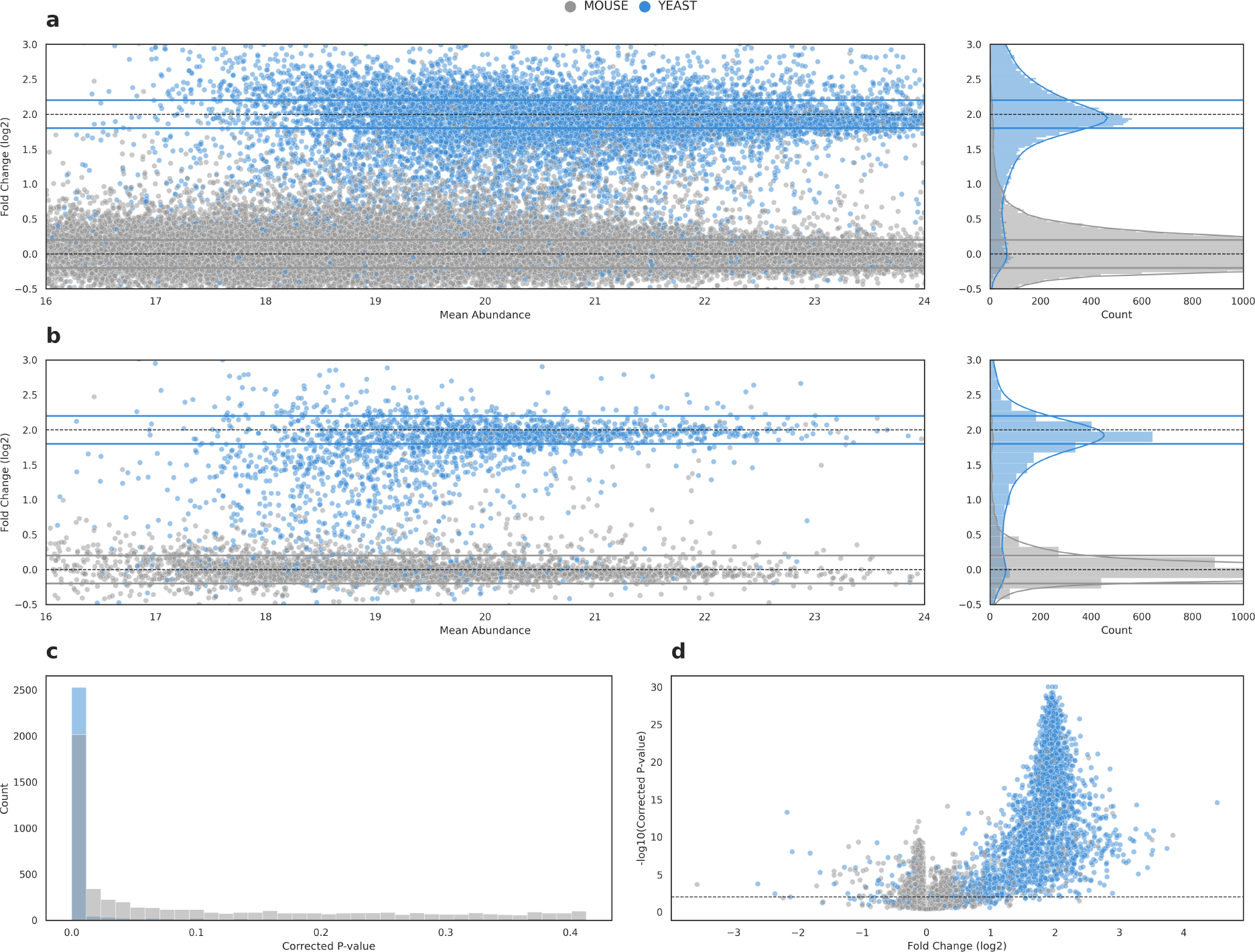
The quantification of proteins from precursors and their differential abundance analysis. The quantification of proteins from precursors can greatly reduce noise in a data set while providing more meaningful annotations for further downstream analysis. **a** shows the distribution of mouse and yeast precursors across the Fold Change (log2) and Mean Abundance dimensions. This visualizes how the quantified precursors group around their expected ratios, indicated by the dotted lines on the scatter and histogram plots. **b** shows the same as **a**, but after the relative quantification of proteins from their precursors. **c** shows the p-value histogram for the statistical comparison between dilution groups. The left skew of the data indicates an easily detectable effect. **d** shows the distribution of Fold Change (log2) compared to corrected p-values in a volcano plot shows the expected fold change of yeast proteins around 4X (2 log2).

Differential expression analysis is a crucial step in the identification of proteins of interest for any biological question. Substantial research has been done on the subject, resulting in different algorithms that all accept data in different formats [6, 7, 9, 10]. With DPKS we make the most common algorithms available in our modular pipeline through our expressive API and Python package, making it easy to select which method you would like to apply to your data (t-test, linear regression, ANOVA) in combination with multiple testing correction. To reduce the impact of missing values, DPKS supports uniform range and uniform percentile imputation algorithms to maximize protein comparisons. To benchmark our differential abundance methods in DPKS, the above 2 selected groups used for protein quantification (4X vs. 16X) from the spike-in data were compared to each other. The distribution of p-values is skewed to the left (**Figure 3c**), indicating that there is a significant difference between the 2 analyzed groups [17]. Additionally, the majority of differentially abundant proteins are yeast proteins with a peak around the expected 4-fold increase mark (a log2 fold change of 2) (**Figure 3d**). All of the described methods above can be easily combined using DPKS and performed in a standardized way. This streamlined, yet flexible, approach encourages reproducibility for classic proteomics differential abundance analysis.

### 2.2 Explainable machine learning uncovers biological signal between proteome states

In additional to classic statistical analysis, it can be useful to develop predictive models that identify proteome states from quantified proteins. This type of analysis could help develop tools for early detection of disease, stratification and characterization of subphenotypes, and biomarker discovery for example. Typically, building and training predictive machine learning models requires expert domain knowledge and programming to complete. With DPKS, we have abstracted away the training of models to a few straight forward function calls, that any experimentalist with basic programming knowledge could use (**Figure 1b**). DPKS provides robust training and evaluation methods, utilizing k-fold cross validation to support experiments of varying sample size, and to provide a distinction between training and testing data for validation. Additionally, methods for hyper-parameter optimization have been provided through randomized search or a novel genetic algorithm to automatically select the best performing set of parameters for the selected classification algorithm (**Supp. Figure 6**). Each of these function calls return the trained model, data scaler, and evaluation results as distinct objects so that they can easily be used outside the DPKS architecture. In order to provide interpretable models that can be analyzed for biological insight, DPKS provides a means to interpret trained models and calculate the importance of each protein in the model using SHAP [14]. The calculated importance of each protein towards a models prediction allows for the data driven selection of the most important features for a particular proteome state instead of using p-value thresholds. As we have previously described [12, 13], proteins found as most important for the classification of a proteome state may not be the most statistically significant, and those with low fold-change may also be important predictive drivers (**Supp. Figure7a**). Finally, DPKS supports the prediction of new data using a trained and interpreted model with a single function call and returns a fully annotated DPKS quantitative matrix to be used for further analysis (**Figure 1b**).

Through the interpretation of trained predictive models, it is possible to gain insight into the proteins that are most important in driving classification of a proteome state that standard differential abundance may miss, but thresholds will still need to be applied to select which proteins to investigate further. To avoid thresholding on protein importance, we have developed a novel protein ranking algorithm that combines recursive feature elimination with XAI via SHAP [14] termed RFE-SHAP (first introduced in Scott et al. 2023[12]). Using quantified proteins from the above described steps as input, we rank all proteins in our dataset through the iterative training and interpretation of consecutive XGBoost [18] classification models in a loop using recursive feature elimination. During training, the worst performing protein is removed per loop and assigned a rank based on the number of proteins considered, until the rank of all proteins has been determined. The classification accuracy at each step is measured so that the set of proteins with the highest accuracy can easily be selected (**Figure 4a**). To evaluate RFE-SHAP, we downloaded a dataset of 687 samples of patients suffering from COVID-19 [15]. Patients from levels 6-7 on the COVID-19 severity scale developed by the World Health Organization (WHO) were considered “severe” as they needed mechanical intervention for respiration, while those *<* 6 were considered “less severe” and did not need mechanical intervention. Based on the complete data set, 406 patient samples were classified “severe” and 281 “less severe”, making up the positive and negative labels in our training data respectively. First, we ranked all proteins in the data set, and identified a panel that maximized accuracy while minimizing the number of proteins used (**Figure 4a**). Throughout the ranking process, predictive accuracy changes as the least important protein is eliminated from the feature space at each training iteration. After all proteins have been ranked, we find the panel of most important proteins by maximizing their predictive accuracy using k-fold (k=3) cross-validation while minimizing the number of proteins used as features. We found a group of 10 proteins (Polymeric immunoglobulin recepto (PIGR), Apolipoprotein F (APOF), Coagulation factor IX (FA9), Complement C3 (CO3), Vitronectin (VTNC), Platelet basic protein (CXCL7), Gelsolin (GELS), Protein AMBP (AMBP), Zinc-alpha-2-glycoprotein (ZA2G), and Protein S100-A11 (S10AB)) to provide the highest accuracy (0.897) when differentiating between severity subphenotypes (**Figure 4c**). The distributions of each individual protein and their relationship to the other proteins in the panel is visualized in **Supp. Figure 8**.

**Figure 4:**
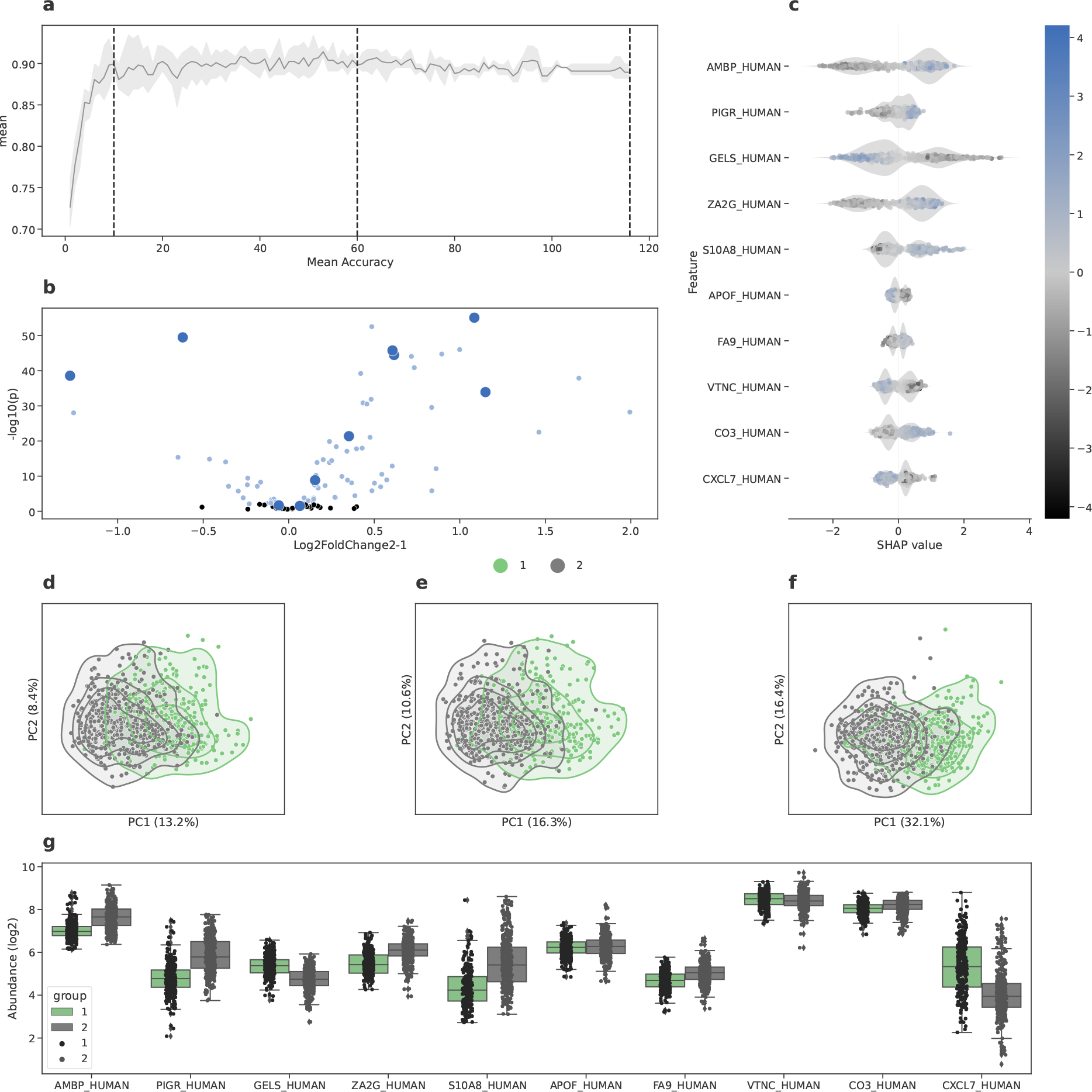
Machine learning analysis and protein panel selection. Machine learning can be used to train highly accurate classifiers and explainable machine learning can be used to select highly accurate panels of proteins that are useful in describing a particular proteome state, in this case severe COVID-19. **a** shows the change in accuracy during feature selection for each iteration of classification as proteins are removed from model training. The dotted lines indicate 3 points during selection that are visualized in **d, e**, and **f. b** shows a volcano plot for the comparison of severe COVID-19 groups. The large points on the plot indicate where the 10 machine learning selected proteins fall on the volcano plot. **c** shows the SHAP distributions for each of the 10 selected proteins. **d, e, f** show the change in PCA from 116 used proteins, 60 used proteins, and then the final 10 selected proteins. **g** shows the quantitative distributions of the selected proteins between groups.

As we have previously described [12, 13], this panel of proteins would have been difficult to identify using classic differential abundance analysis, and emphasizes the importance of integrating machine learning methods into existing analysis (**Figure 4b**). Interestingly, some of the important features do not have significant p-values, and some of the more significant (low p-value) proteins are not considered most important in differentiating between severity subphenotypes of COVID-19. In our previous analysis of the COVID-19 data [13], we selected a threshold of the top 10 most important proteins for classification using BINN, while here we optimize and automate the selection of the most predictive panel of proteins as a group using RFE-SHAP. Here, the most important proteins (low feature rank) correspond to initial high SHAP values, although some proteins initially considered important are removed as the feature space gets smaller (**Supp. Figure7b**). Additionally, many proteins with significant p-values are not considered important in classification and removed early from the model. While converging on the optimal set of proteins used for classification, the separation between subphenotypes using PCA increases when the most accurate panel of proteins is applied (**Figure 4d-f**). The abundance profiles for each of the 10 proteins are visualized in **Figure 4g**. Additionally, we demonstrate how this panel of 10 proteins can effectively separate the COVID-19 severity subphenotypes using hierarchical clustering with the 10 selected proteins (**Figure 5a**), where accuracy was evaluated using the rand index (0.75).

**Figure 5:**
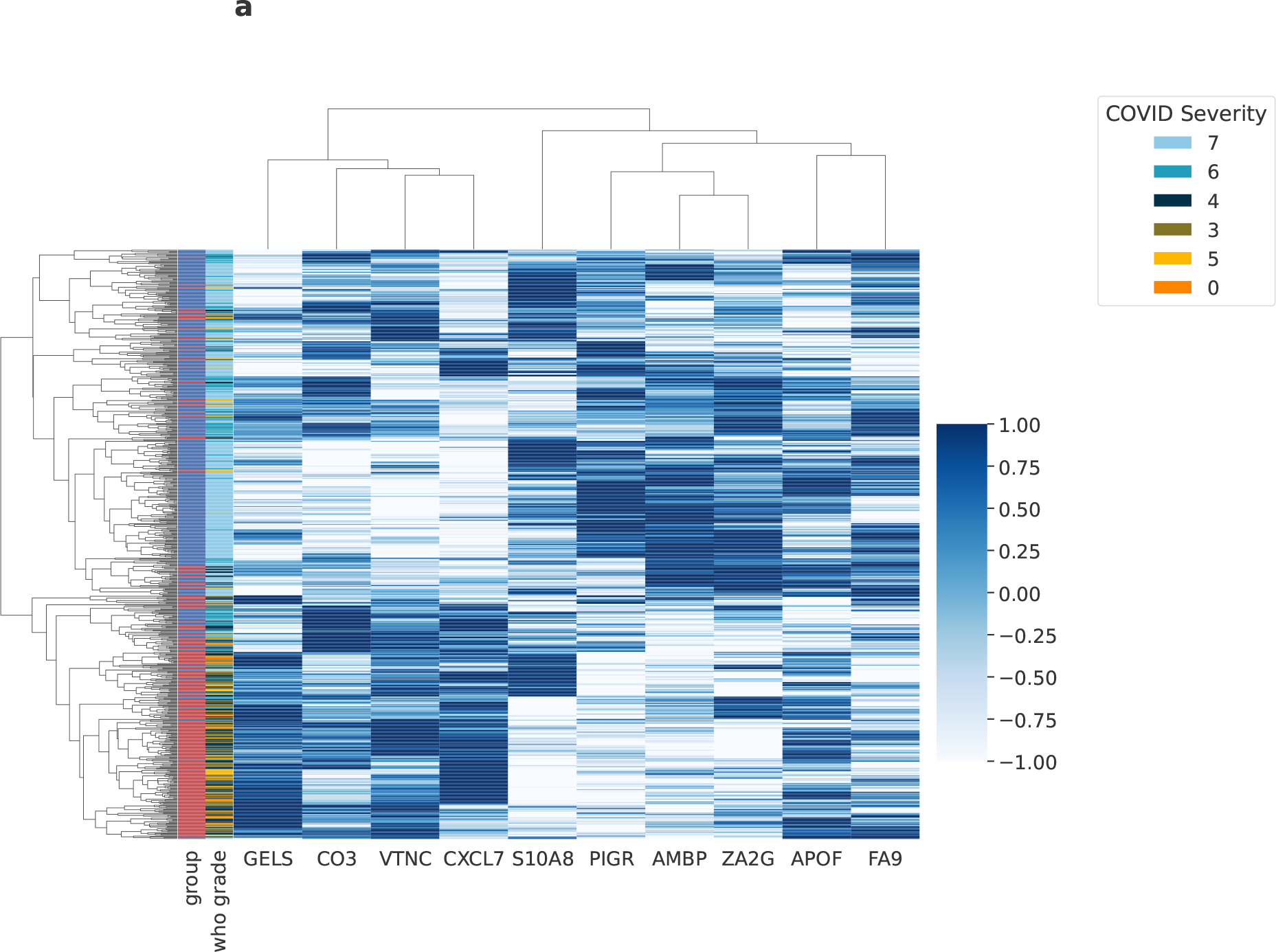
Cluster map using the panel of selected proteins. The clustermap shows how the groups of COVID-19 severity cluster using the 10 selected proteins from the RFE-SHAP analysis above. These 10 proteins accurately stratify COVID-19 severity subphenotypes and provide a set of proteins for further investigation. There are 2 main clusters of proteins, those that are up in severe COVID-19 and those that are down in severe COVID-19.

As with the above statistical analysis, feature ranking via RFE-SHAP is easily available from the DPKS API using just 1 line of code (**Figure 1b**), exposing complicated methods to those without machine learning expertise. Using the generalized approach available with the DPKS library, researchers can easily apply these methods to their own biological systems to train predictive models and identify the important proteins for whatever proteome is being investigated without any thresholding.

## 3 Discussion

With a focus on usability and speed, DPKS provides a suite of high-performance algorithms that allow researchers to focus on answering biological questions instead of getting lost in analytical details. As there are so many different tools for each analytical step, the task of selecting which to use for an analysis can be overwhelming. DPKS makes these decisions easier by providing a framework for best practices that are easily applied to raw quantitative data with just a few lines of code. If certain default options need to be changed, it is trivial to customize functional steps for specific analytical needs. Although there may be other packages, written in both R and Python, that provide access to normalization, protein quantification, and differential abundance analysis, they all require differently formatted data, so piping the output of one tool into another may not be so straightforward. Additionally, no available packages expose step-by-step functionality from raw quantitative data to a highly predictive machine learning model and fully ranked features for protein selection with minimal code.

The API of DPKS was designed with the end user in mind and attempts to provide consistent access to functionality, backed by documented and well structured modular code. Basic object oriented principles, with minimal abstraction, keep the code that powers the available algorithms clean and concise. The back-end functionality of DPKS is split into functional modules that are exposed to the user via a common data structure. These consistent design principles makes the core functionality easily extensible and approachable for open-source developers as new functionality can easily be added without breaking other aspects of the code. The focus on simple clean code means that some state-of-the-art methods may not be available in DPKS without a native Python implementation in the library. Fortunately, it is also possible to export tabular data at any time to perform computation outside of DPKS and easily import the altered quantitative data back into DPKS for further analysis.

Recently, a variety of feature attribution methods have been introduced [14, 19, 20] that allow for the interpretation of complex machine learning models. In the case of DPKS we have implemented a novel feature ranking algorithm using SHAP [14] and recursive feature elimination that provides a rank to each individual feature in a dataset along with numerical values for the importance of each feature. The benefit of ranking each individual feature allows for the granular investigation of the predictive power of each protein in a system and to ensure that the most accurate panel of proteins possible are selected. Due to the inherent flexibility of DPKS, it would be simple to implement additional feature attribution methods for feature ranking if an alternative method wanted to be used. Although we specifically use RFE-SHAP in this study to select important proteins, proteins can further be filtered after differential abundance analysis to only include significant proteins below a certain p-value threshold as an effective means of feature reduction due to the flexible and modular nature of DPKS.

One main benefit of DPKS is the expressive and flexible API that exposes complex algorithms to users that may not have expert programming knowledge. This is especially relevant in the *classification* module of DPKS which allows users without machine learning expertise to train highly predictive classifiers and apply feature attribution methods to identify the most important features in a system. This means that researchers can easily apply methods previously available to only expert machine learning practitioners and allow them to focus on the biological interpretation of their results. The flexible nature of the library also allows machine learning experts to apply their own models, as well as manipulate the models trained using DPKS, enabling custom analysis and pipelines to be easily created. Additionally, due to extensive optimization, the user friendly API of DPKS is backed by highly performant algorithms to allow for efficient experimentation of the available methods in the package if needed for a particular analysis.

With DPKS, we present the first full featured analytical platform in the Python language to go from a raw quantitative matrix to highly accurate explainable predictive models and biomarker panels in minutes.

## 4 Methods

### 4.1 DPKS

DPKS is implemented in Python and is open source and freely available under an MIT license (https://github.com/InfectionMedicineProteomics/DPKS). All implementation details for the methods descrived below are available in the source code.

#### 4.1.1 Parsing

DPKS provides parsers for the output for a variety of label free quantification software (DIA-NN [21]) or a general input file based on the output from GPS [12]. This allows for the output of any quantification to be used as input to DPKS.

#### 4.1.2 Normalization

The normalization methods in DPKS are designed to help reduce technical variation between samples. For example, for the mean normalization method in DPKS, we first calculate the means for each sample represented by vector *S*_*m*_ = [*S*_1_, *S*_2_, *S*_3_, …*S*_*n*_], and then the mean of *S*_*m*_ as the adjusting factor *S*_*tot*_. Finally, each sample in the quantitative matrix *X* is divided by their mean, multiplied by the adjusting factor, and logarithmized to give the normalized matrix.

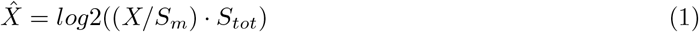

Depending on the method used, the calculation of the sample factors will be performed differently. DPKS currently implements normalization methods for mean, median, and total ion chromatogram normalization methods.

Additionally, DPKS exposes a retention time sliding window normalization method based on the implementation in the NormalyzerDE R package [1]. Here, data in the quantitative matrix is normalized in slices of overlapping retention time to more accurately correct specific effects during different time points in the gradient. Precursors in the matrix are first sorted by their chromatographic retention time and placed in bins of a certain minimum data points. These retention time bins are then iterated and normalized individually to produce the final quantitative matrix. The use of overlapping windows can be specified if desired.

#### 4.1.3 Quantification

Quantification can be done at the precursor, peptide, or protein level in DPKS. To provide a method of absolute quantification, the TopN method selects the top-n number of features for a given summarization level and combines them with a particular function (sum, mean, or median). Relative quantification is available through an implementation of the iq algorithm [3], where the optimal ratios between given feature at the summarization level are used to adjust quantities and provide an overall relative abundance of an analyte compared to the different samples. In order to provide rapid results, this algorithm is optimized using numba and just-in-time (JIT) compilation to compile native Python to machine code for faster analysis. This method scales well with increasing numbers of samples as many of the computations are vectorized, but run time is increased as the number of features quantified increases.

#### 4.1.4 Differential abundance

Differential abundance analysis can also be performed at the precursor, peptide, or protein level in DPKS. Based on the indicated level, DPKS will iterate through the quantified features and compare them between groups indicated in the design matrix using one of the available statistical tests (t-test, linear regression, or ANOVA) followed by multiple testing correction for false discovery rate control. The DEScore, first introduced in Hartman et al. [13], can be used to select out features that are the most differentially abundant as an alternative to selecting the lowest corrected p-values.

#### 4.1.5 Imputation

To fill in missing values for differential abundance analysis or machine learning, DPKS provides easy programmatic access to 2 main imputation methods. Uniform range imputation allows the user to provide a range of values which a uniform distribution is sampled from, while uniform percentile imputation allows the user to indicate a percentile of the data to sample a uniform distribution from.

#### 4.1.6 Machine learning

DPKS provides access to a variety of machine learning techniques for building and interpreting classifiers. From the base quantitative matrix data structure in DPKS it is possible to train, optimize, evaluate, and apply any classification algorithm that adheres to the scikit-learn API. Training models using the DPKS interface also provides direct integration with explainable artificial intelligence and feature attribution methods via SHAP [14]. All model training is performed using k-fold cross validation for robust model evaluation. After training, you can easily interpret the trained model to calculate the importance of each individual precursor, peptide, or protein towards the classification of individual samples. Additionally, it is possible to rank all proteins in a quantitative matrix to find the proteins that maximize the accuracy of classification between the groups being analyzed. This ranking is done using recursive feature elimination (RFE) with SHAP as a means of calculating feature importance. Using k-fold cross validation, the importance of each feature is calculated and the least important features are pruned from the list used to train the model. The protein rank is assigned as the number of features left in contention before the removal of the current worst feature. From here, a group of proteins can be identified that maximize accuracy in quantification. The model trained from this subset of proteins can then be easily applied to new data using the DPKS API. Model hyper-parameter optimization is available via the scikit-learn implementations of random and grid search, as well as a third method using genetic algorithms to select the best populations of hyperparameters for a given model and feature space.

#### 4.1.7 COVID-19 data analysis

COVID-19 data was downloaded from PRIDE [22] using the accession code PXD025752[15]. Patients from levels 6-7 on the COVID-19 severity scale developed by the World Health Organization (WHO) were considered “severe” as they needed mechanical intervention for respiration, while those *<* 6 were considered “less severe” and did not need mechanical intervention. Raw precursor data was normalized using retention time window sliding normalization with mean as the applied function. Proteins were quantified using the iq [3] implementation in DPKS. Differential abundance analysis was performed using linear regression models, and multiple testing correction was performed. Immunoglobulin and hemoglobin proteins were removed from the quantitative matrix prior to applying the machine learning functionality of DPKS. RFE-SHAP was performed and the group of proteins that maximized accuracy while minimizing the number of proteins used was selected as the most predictive usable panel.

## 5 Acknowledgements

LM was supported by the Swedish research council (grant number VR-2020-02419), the Wallenberg foundation (grant number 2016.0023) and Alfred Österlunds Foundation. JM was supported by the Wallenberg foundation (WAF grant number 2017.0271), the Swedish research council (grant number 2019-01646 and 2018-05795) and Alfred Österlunds Foundation.

## 6 Author contributions

AS and LM conceptualized DPKS. AS, LM, and EH wrote code and implemented the algorithms within DPKS. AS wrote the manuscript and designed the study. JM provided supervision and contributed to the writing of the manuscript. LM oversaw and supervised the overall project and contributed to the writing of the manuscript. All authors approved the manuscript.

## 8 Data availability

The spike-in data was downloaded from the ProteomeXchange Consortium via the PRIDE [22] partner repository with the dataset identifier PXD038377.

## 9 Code availability

All DPKS code is open-source and freely available under the MIT license at https://github.com/ InfectionMedicineProteomics/DPKS.

## 10 Inclusions & Ethics

## 11 Supplement

**Figure 6:**
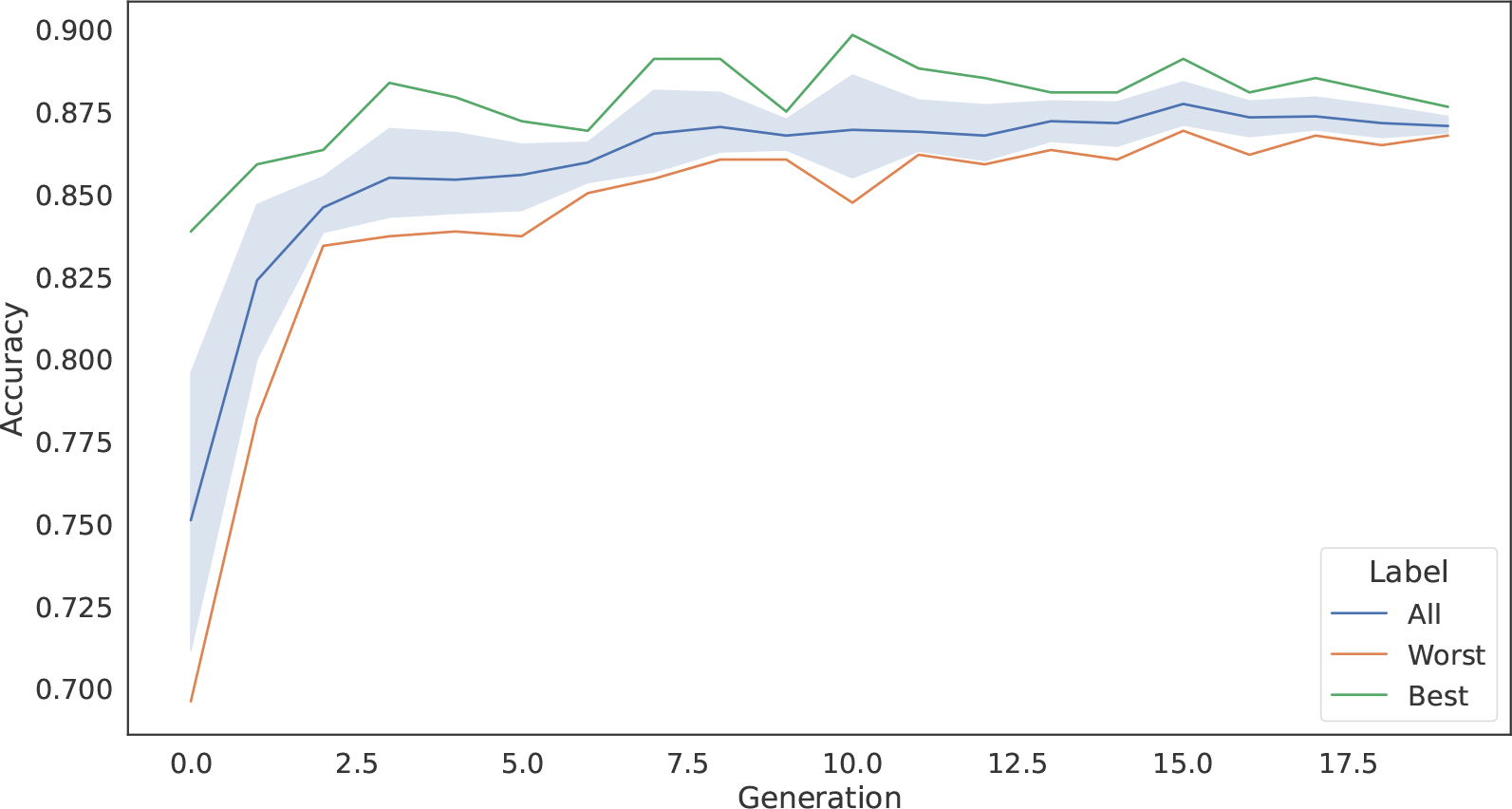
Optimizing parameters. This plot shows how the accuracy of a classifier increases as hyperparameters are tuned using a genetic algorithm. As the generations progress, the accuracy of a model converges on a set of optimal hyperparameters.

**Figure 7:**
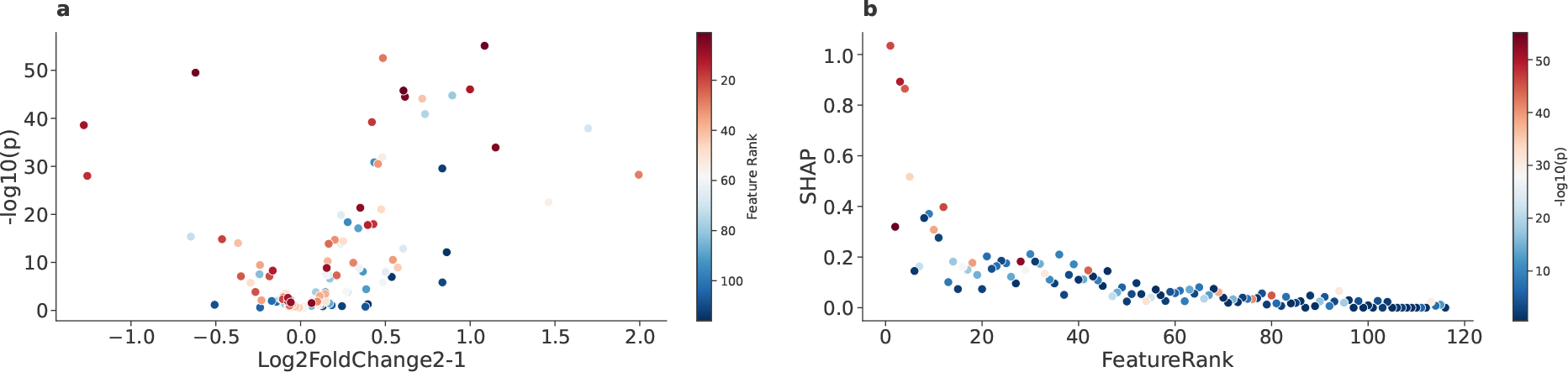
Feature rank scatter plots. These plots show the relationship of feature importance in a classifier against standard differential abundance and feature rank. **a** is a volcano plot that visualizes log2 fold changes against adjusted p-values and colors them by feature importance (SHAP value). Many of the most important features for calssification are not the proteins with the highest fold-change or the lowest p-values. **b** visulizes how the rank of features can change as features are removed from consideration during feature selection, although the most important features usually remain the highest ranked.

**Figure 8:**
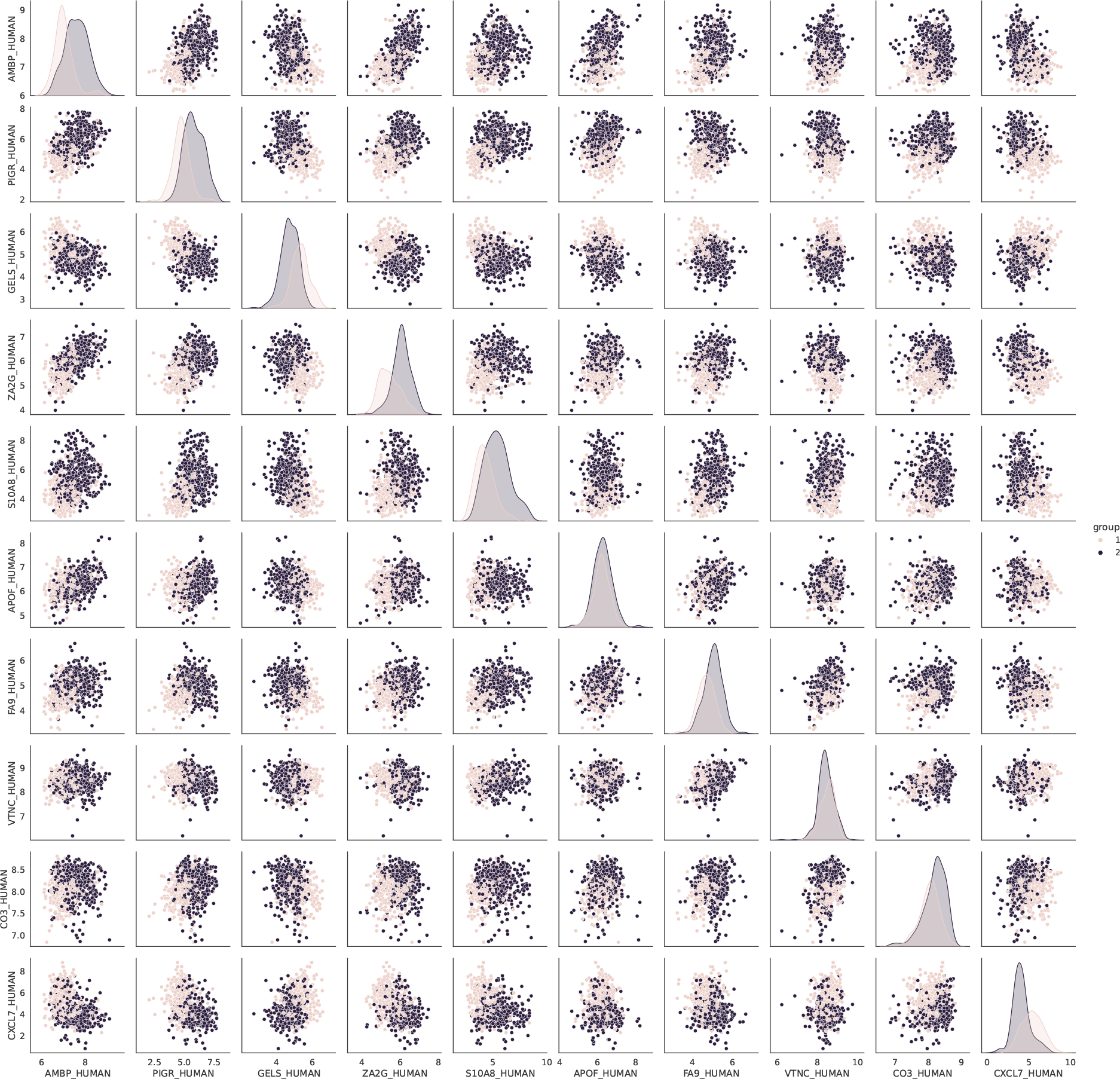
Feature relationship plots. All selected proteins and their relationship to all other proteins in the selected panel are visualized here. Some proteins that do not have a significant difference between labels are highly effective at separating classes when combined with other features.

